# SpaceBlocks: an expert-supervised workflow for cohort-level spatial transcriptomics analysis

**DOI:** 10.64898/2026.10.30.763100

**Authors:** Sergio Oterino-Sogo, Nazim Mechkouri, Daniel García-Ruano, Alexis Groppi, Hamid-Reza Rezvani, Macha Nikolski

## Abstract

**Summary:** Spatial transcriptomics (ST) has become a central tool in tissue biology, with multiple technologies addressing the issue from a different angle. Each platform imposes its own raw data format conventions, and few workflows address reproducible, scalable, cohort-level analyses while integrating expert-supervised decisions to strengthen computational and biological provenance.

In this work, we present SpaceBlocks, a Snakemake workflow for reproducible cohort-level analysis of single-cell resolution-ST data that prioritizes expert supervision over full automation. SpaceBlocks homogenizes heterogeneous platform outputs into a common data format, whose structure is validated to ensure downstream processing. Comprehensive expert supervision is facilitated through per sample quality control and cell type annotation diagnostic visualizations, after which downstream analyses automatically follow. SpaceBlocks also allows the ingestion of technology-agnostic externally formatted data and provides interoperability with existing tools and technologies.

We demonstrate the use of SpaceBlocks through the reanalysis of 37 samples from public Visium HD, Xenium 5K and MERSCOPE datasets. SpaceBlocks exhibited processing scalability across the three platforms. This reanalysis identified previously missed cell types in Visium HD and heterogeneous marker gene expression in MERSCOPE, highlighting the need for expert-supervised annotations. Moreover, SpaceBlocks facilitated the identification of differentially expressed genes associated with previously defined multiple sclerosis (MS) spatial domains in MERSCOPE, and transcriptional differences between healthy and malignant colorectal tissues in Visium HD. Together, these findings establish SpaceBlocks as a flexible and reproducible workflow for expert-supervised spatial transcriptomics analysis across technologies and biological cohorts.

**Availability and Implementation:** SpaceBlocks is freely available under the MIT license on GitHub https://github.com/cbib/SpaceBlocks and the official Snakemake catalog https://snakemake.github.io/snakemake-workflow-catalog/docs/workflows/cbib/SpaceBlocks.html. SpaceBlocks v0.5.1 code, config files, and the public data formatting scripts used to reproduce the results from this study have been deposited in Zenodo (doi: 10.5281/zenodo.23378236).

**Supplementary information:** Supplementary information is available online. Conflict of interest: None declared.

## 1. Introduction

Spatial transcriptomics (ST) was recognized as the Method of the Year in 2020 (Marx, 2021). Over the past decade, spatial sequencing- and imaging-based technologies have expanded rapidly (Cervilla, et al., 2026; Lim, et al., 2025; You, et al., 2024), enabling increasingly detailed characterization of tissue organization while introducing heterogeneous conventions for count matrices, spatial coordinates, images and annotations. A diverse software ecosystem has consequently emerged, supporting segmentation, quality control (QC), normalization, clustering, annotation and spatial analysis (Marco Salas, et al., 2025; Singhal, et al., 2024; Specque, et al., 2025). Now, several frameworks exist, trying to address complementary needs: Squidpy and Giotto Suite offer extensible analysis frameworks for general analyses in Python and R (Chen, et al., 2025; Palla, et al., 2022), nf-core/spatialvi automates the processing of 10x Genomics Visium data (Ewels, et al., 2020), and Sopa provides a technology-invariant pipeline for image-based spatial omics (Blampey, et al., 2024).

These ecosystems provide rich analytical functionality, but reproducible, cohort-level orchestration of the full sequence of expert-supervised decisions, starting from heterogeneous platform outputs remains a challenge. During analysis, expert supervision is required through QC, cell type and region of interest (ROI) annotation, and the definition of the experimental design for statistical contrasts, ultimately leading to interpretable results. In practice, these decisions are made in notebooks or ad hoc scripts and rarely survive publication with enough detail for full reproducibility (Ziemann, et al., 2023). Thus, there is a critical need for workflows that systematically capture analytical decisions as explicit, versioned inputs, automate routine processing steps, and formally integrate expert judgment to ensure transparency, consistency, and reproducibility.

Here, we present SpaceBlocks, an expert-supervised Snakemake workflow for reproducible and scalable cohort-level analysis of single-cell resolution ST data. SpaceBlocks centers on a standardized AnnData contract format that provides a single validated entry point to all downstream analyses. The workflow accounts for two subworkflows. Technology-specific HeadBlocks transform raw platform outputs into the standardized AnnData contract format, whereas a decoupled mode accepts externally prepared datasets satisfying this specification. Following validation, a shared CoreBlocks subworkflow performs the full downstream analysis identically across platforms, including QC, clustering, cell type annotation, integration, spatial neighbourhood and domain analysis, and reporting. To identify differentially expressed genes, SpaceBlocks aggregates raw counts per sample into pseudobulk profiles across all cells, optionally stratified by cell type and/or tissue region. Sample- and region-level metadata then define the comparison groups, covariates, and pairing structure used by DESeq2. DESeq2 supports user-defined paired or unpaired Wald contrasts, additive categorical covariates, optional one-vs-rest tests, and an omnibus likelihood-ratio test for multilevel experimental designs. SpaceBlocks connects specialized algorithms and expert judgement through a standardized and auditable workflow that can run both locally and on compute clusters, promoting transparency, reproducibility, and scalability for single cell resolution ST analyses. Its modular architecture and shared validation entry point lets the workflow be extended to new technologies and new downstream analyses without requiring modifications to the core downstream workflow.

## 2. Implementation

SpaceBlocks was developed as a Snakemake (≥ v9.13) workflow organized into two subworkflows, the HeadBlocks and the CoreBlocks. Necessary software is installed by Snakemake using Conda and the supplied YAML environment configuration, specified in each of the Snakemake rules. All Snakemake rules use Python (v3.10) dependencies, except for the differential expression testing, which runs in R (v4.3) using the native DESeq2 (≥ v1.4.0) implementation. SpaceBlocks currently supports raw outputs from MERSCOPE, Visium HD, Xenium 5K and Atera. Atera support from the HeadBlocks is provided as a preview, since it was developed from the pre-release datasets, and the platform’s commercial release may slightly modify the instrument’s raw file structure. Data from other spatial transcriptomics platforms can be preprocessed and incorporated to the workflow by complying with the standardized AnnData data contract. The modular architecture also enables future extension of HeadBlocks to support additional technologies directly from their raw outputs.

### 2.1. Workflow architecture and data contract

The architecture of SpaceBlocks is shown in Figure 1a, and schematic outputs are shown in Figure 1b. Each run is specified through a schema-validated YAML configuration file and tab-delimited sample sheets, which centralize QC, normalization, spatial neighbourhood analysis, clustering and UMAP parameters, alongside sample level experimental data and metadata.

**Figure 1.**
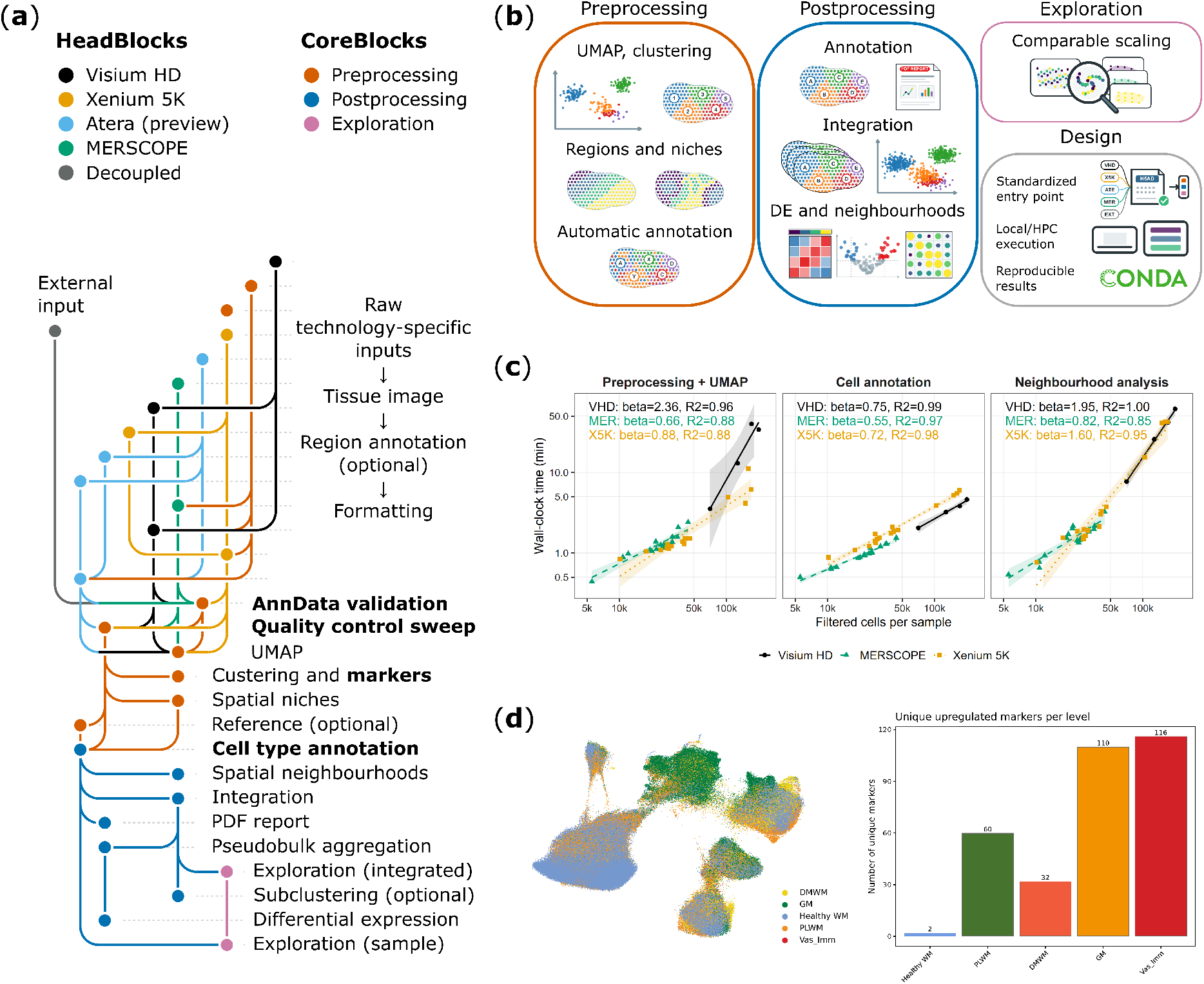
Overview of the SpaceBlocks workflow. (a) Snakevision tubemap scheme of the workflow rules, coloured by technology and processing step. SpaceBlocks v0.5.1 supports Visium HD, Xenium 5K, MERSCOPE and Atera (preview) processing from their raw platform outputs. The AnnData contract validation entry point and expert-supervised steps are displayed in bold font. (b) Schematic overview of the outputs produced by the CoreBlocks subworkflow and the main design features of the pipeline. (c) Runtime benchmarking across Visium HD (n = 4), Xenium 5K (n = 16) and MERSCOPE (n = 17) datasets. Associations between runtime and dataset size were evaluated using separate ordinary least-squares regressions of log10 wall-clock time against log10 filtered cell number, where each sample represents a single data point. β denotes the fitted scaling exponent, where doubling the cell count multiplies the estimated wall-clock time by 2^β^. (d) Application of SpaceBlocks to the multiple sclerosis MERSCOPE dataset published by Feng et al., 2025. Left, the originally published BANKSY niches projected in the harmony-integrated UMAP generated by SpaceBlocks. Right, the number of domain-specific upregulated genes from one-vs-rest tests (Benjamini-Hochberg p-adj < 0.05 and | log2FC| > 0.25).

The workflow accepts platform-specific raw data formats as input, and provides a decoupled execution mode that bypasses the HeadBlocks to accommodate externally preformatted datasets. Rule-associated Conda environments isolate software dependencies between Snakemake rules. The workflow ships with execution profiles for local runs and for SLURM clusters, each setting per-rule resources, retry policy, logging and Snakemake benchmarks. Additional schedulers can be targeted by adding the corresponding Snakemake profile.

Each HeadBlock produces a single unfiltered contract file per sample, specifically an AnnData H5AD file (Virshup, et al., 2024). This contract contains non-negative integer expression counts, unique cell identifiers, sample information, and finite spatial coordinates, with tissue images and associated scaling information included when available. A dedicated Snakemake validation rule verifies compliance with these requirements before downstream processing begins. Contract violations, such as non-integer counts and/or missing spatial coordinates abort the workflow before any analysis runs. Optional elements (tissue image, region annotation, mitochondrial genes) are recorded as warnings, and downstream rules consequently adapt (for example, spatial plots fall back to a coordinate scatter when no image is present) so the execution continues.

### 2.2. Reproducible expert-guided analysis

SpaceBlocks is semi-automated by design, with QC and cell type annotation implemented as explicitly supervised checkpoints, informed by automated tools.

Before filtering, an optional QC sweep evaluates a range of common technical thresholds per sample, visualized through violin, scatter and spatial plots to inform expert decision-making. Once QC thresholds are set in the configuration, data is log normalized, and Leiden clustering is evaluated across a range of resolutions. UMAP and spatial cluster maps, gene expression plots, silhouette scores and pyclustree (v0.4) visualizations (Zappia and Oshlack, 2018) support the selection of an appropriate clustering resolution.

### 2.3. Cohort-level downstream analysis

To inform cell type annotation, SpaceBlocks accepts externally annotated single cell references (H5AD) and curated marker gene tables (TSV). Cell type annotation may be supplied as an expert-curated mapping between clusters and cell types in TSV format, transferred from an annotated single-cell reference using *scanpy*.*ingest*, or imported as externally generated cell-level labels. Optionally, histological region annotations may be supplied using GeoJSON stored polygons, which can be easily generated by QuPath (≥ v0.5.1) (Bankhead, et al., 2017).

The workflow incorporates spatial analyses to characterize cellular and tissue organization at both sample and cohort levels. Squidpy-based neighbourhood enrichment and co-occurrence analyses quantify spatial relationships between annotated cell types within samples and tissue regions (Palla, et al., 2022). To identify shared spatial patterns across the cohort, individual samples are concatenated into a single AnnData object. Within this joint object, BANKSY identifies spatial niches by offsetting sample coordinates, so neighbourhoods remain within sample, producing a common niche representation that is then mapped back to each tissue (Singhal, et al., 2024).

SpaceBlocks applies Harmony batch correction to a configurable metadata variable (Korsunsky, et al., 2019) and generates a geometric sketch for scalable visualization (Hie, et al., 2019). For differential expression, pseudobulk raw count matrices are summed by biological sample using the decoupler Python package and optionally stratified by cell type and/or tissue region (Badia, et al., 2022). SpaceBlocks supports user-defined configuration of paired or unpaired Wald contrasts by cell type, tissue region, metadata variables, or a combination of them. In unpaired designs, the biological sample variable (specified in the sample sheet) is used as the experimental unit. On the other hand, paired designs treat samples as the matching variable, and only complete biological pairs are retained for analysis. As a safeguard, repeated within-sample observations are rejected in unpaired analyses, unless the user explicitly accepts the assumption that they are independent. For multilevel experimental designs, the workflow also allows one-vs-rest Wald tests and summarizes unique findings, enabling condition-specific exploratory biomarker discovery. SpaceBlocks also supports an omnibus likelihood ratio test for multilevel experimental designs, with significant genes subsequently clustered by expression pattern using the DEGreport R package (≥ v1.46). Minimum number of cells, counts, and replicate thresholds are configurable and applied before testing, and if any contrast is not possible, the reason is reported. All comparisons are performed with DESeq2 (≥ v1.4.0) original implementation in R (Love, et al., 2014). Outputs include complete tables, volcano plots, metadata-annotated heatmaps and bar plots, and gene-expression patterns plots for likelihood-ratio test results.

### 2.4. Extended exploratory analyses

An optional subclustering rule re-clusters chosen annotated population/s (e.g. neurons) to resolve finer subtypes. Two complementary exploration rules then plot user-specified genes and gene sets (one on the integrated cohort object, one per sample) using a shared expression scale across both views so that intensities are directly comparable across samples.

The workflow records intermediate and final AnnData objects, tabular results, diagnostic figures, logs and benchmarks into a standardized output hierarchy. A per-sample PDF report summarises QC, clustering, cell type annotations, tissue regions, spatial niches and cell type composition. These reports complement the AnnData objects and tabular outputs, providing an accessible overview of the analytical results while preserving the underlying data for downstream exploration and reuse.

## 3. Application

### 3.1. Platform coverage and computational performance

We illustrate SpaceBlocks processing time for 17 MERSCOPE samples containing 5,550-44,000 filtered cells (Feng, et al., 2025), 4 Visium HD samples containing 70,618-201,093 filtered cells (10x Genomics, 2025) and 16 Xenium 5K samples containing 10,167-171,526 filtered cells (Dumoulin, et al., 2026) (Figure 1c, see Data Availability).

On the benchmark system (Rocky Linux 9.4 with SLURM scheduler, HPE Apollo 2000 Gen10+, AMD EPYC 7513 CPUs, PC4-3200AA RAM), the median per-sample wall time for shared CoreBlocks rules was 7.35 min for MERSCOPE (3.01-12.80 min range), 101.96 min for Visium HD (29.02-156.88 min range) and 9.48 min for Xenium 5K (4.46-84.31 min range). Extended wall times and statistics by rule and technology are shown in Supplementary Figure 1 and Supplementary Tables 1-2. Wall-clock time was positively associated with the number of filtered cells across the three platforms. UMAP runtime differed systematically between platforms, potentially reflecting differences in the number of features profiled (500 for MERSCOPE, 5,000 for Xenium 5K, 20,000 for Visium HD; Figure 1c).

### 3.2. Real-data use cases

#### 3.2.1. Multiple sclerosis profiled with MERSCOPE

We illustrate the use of SpaceBlocks on the MERSCOPE dataset from Feng et al. 2025, which studied multiple sclerosis (MS) progression. The study contains 17 MERSCOPE samples of patients with healthy white matter (n = 3) and lesional MS tissue (n = 14), with samples annotated as leukocyte-rich (LR) and leukocyte-poor (LP) demyelinated white matter

(DMWM), perilesional white matter (PLWM), and healthy white matter (WM). The study provided annotations for cell populations, including oligodendrocytes, astrocytes, microglia and macrophages, neurons, vascular cells and lymphocytes. The study also provided annotated BANKSY spatial domains, including healthy white matter, grey matter, vascular-immune areas, perilesional and lesional white matter. We used the published cell type annotations to benchmark upstream processing and the BANKSY domain annotations to drive downstream differential expression between MS lesion subtypes.

The data was initially pre-processed to enter the SpaceBlocks workflow skipping the HeadBlocks (Availability and Implementation). In the CoreBlocks, SpaceBlocks QC sweep evaluated the study cut-offs in technical parameters (Supplementary Figure 2), indicating that the GEO accessible dataset was already trimmed for extreme low and high-count observations. The pre-processing step revealed heterogeneity in cell type markers for annotated cell types (Supplementary Figure 3). To further evaluate cell type annotation, we supplied the single nuclei dataset from the same study in SpaceBlocks to perform automatic annotation via *scanpy*.*ingest*. In this specific example, the annotation transferred from the single-nucleus reference dataset using *scanpy*.*ingest* assigned most cells to the oligodendrocyte label, whereas the published cell-type annotations showed a broader distribution of cell types (Supplementary Figure 4). This discrepancy highlights the importance expert supervision when assessing cell type annotation in spatial transcriptomics data.

Finally, we used cell type and BANKSY spatial domain annotations to perform pseudobulk differential gene expression analyses across MS lesion subtypes. SpaceBlocks identified more than 300 differentially expressed domain-associated genes across MS histological lesion subtypes (Figure 1d). The resulting domain-associated signatures mapped to distinct cellular compartments, consistent with the spatial organisation reported by Feng et al. (2025) (Supplementary Figure 5). Notably, none of the signatures was preferentially detected in oligodendrocyte precursor cells (OPCs), a progenitor population central to remyelination. This observation deserves further investigation and illustrates how SpaceBlocks can highlight unexpected biological patterns that require expert interpretation. Taken together, these results emphasize the need for both cell type and domain specific exploratory analyses in ST analyses, which SpaceBlocks facilitates.

#### 3.2.2. Colon cancer profiled with Visium HD

We also used SpaceBlocks to explore the 10x Genomics Visium HD dataset on colorectal cancer, containing 2 healthy and 2 malignant tissue samples without predefined cell type annotations (10x Genomics, 2025). We manually annotated the samples by assigning cell types to the clusters resulting from Leiden clustering (resolution 0.8). Here, SpaceBlocks enabled the identification of 3 cell types that had been missed by the publicly available 10x Genomics notebook (10x Genomics, 2025). Namely, endothelial cells, macrophages and a group of cells with mixed Enteric glial/Schwann-like expression profile (Supplementary Figure 6a). Furthermore, exploratory pseudobulk differential expression analysis identified 1,952 differentially expressed genes between 2 healthy and 2 malignant samples (Supplementary Figure 6c-d). These analyses illustrate the ability of SpaceBlocks to support expert-guided cell type annotation and comparative transcriptomic analyses of spatially resolved tissue samples.

## 4. Conclusions

SpaceBlocks provides a standardized framework for cohort-level ST analysis, built around a validated AnnData contract that separates platform-specific preprocessing from a shared downstream analytical workflow, and formalises the expert decisions such as QC thresholds, clustering resolution, annotation, region definitions, as explicit, versioned inputs. Thus, SpaceBlocks enhances transparency, reproducibility and reusability compared with common ad hoc analyses. Applied to MERSCOPE, Visium HD and Xenium 5K cohorts, SpaceBlocks reproduces the broad published structure while surfacing findings that require further expert evaluation and interpretation (Supplementary Table 3). Its modular, contract-based architecture keeps the workflow extensible to new platforms and interoperable with existing processing tools such as Sopa or ENACT, which can feed pre-formatted datasets through SpaceBlocks’ decoupled mode.

SpaceBlocks v0.5.1 currently relies on vendor cell segmentation outputs, including proprietary Space Ranger for Visium HD. A planned open-source HeadBlock based on STAR-solo and StarDist will enable fully redistributable Apptainer/Docker builds. CosMx and Atera post-release support, and additional normalisation and integration options are under active development.

## Supporting information

Supplementary Table 1. SpaceBlocks per sample wall times across MERSCOPE, Visium HD and Xenium 5K samples.

Supplementary Table 2. Statistics on the per sample wall times linear modelling.

Supplementary Table 3. Comparison between SpaceBlocks and similar existing workflows and tools.

## Acknowledgements

The authors wish to thank Dr. Domitille Chalopin-Fillot and Mr. Mikaël Georges (IBGC, Bordeaux University) for their valuable feedback during the development of SpaceBlocks.

Authors would like to thank Khurasan Studio for granting them full permission of use for their Pavelt font for the SpaceBlocks logo for the scientific publication and GitHub repository.

## Author contributions

Sergio Oterino-Sogo (Conceptualization [lead], Data Curation [lead], Formal Analysis [lead], Methodology [equal], Validation [equal], Writing—original draft [lead], Writing—review & editing [equal]), Nazim Mechkouri (Resources [equal], Methodology [equal], Validation [equal], Writing—review & editing [equal]), Daniel García-Ruano (Resources [equal], Methodology [equal], Writing—review & editing [equal]), Alexis Groppi (Conceptualization [equal], Supervision [equal], Writing—review & editing [equal]), Hamid Reza-Rezvani (Conceptualization [equal], Supervision [equal], Writing—review & editing [equal]), Macha Nikolski (Conceptualization [lead], Supervision [equal], Writing—review & editing [lead]).

## Funding

H.R.R. gratefully acknowledges supports from ITMO Cancer of Aviesan within the framework of the 2021-2030 Cancer Control Strategy, on funds administered by Inserm and Fonds de Dotation pour la recherche de la «Société Française de Dermatologie (SFD)». Computational resources and infrastructure used in present publication were provided by the Bordeaux Bioinformatics Center (CBiB - US Ubx 006). These resources were funded by the CPER DOREMI (Région Nouvelle Aquitaine and Ministère de l’Enseignement Supérieur).

## Public data availability

The MERSCOPE dataset reanalysed in this publication can be accessed at the GEO database (GSE284005). The Visium HD samples were downloaded from the 10x Genomics data portal (10x Genomics, 2025). The Xenium 5K samples were downloaded from Zenodo repository, accession number 19868428 (Dumoulin, et al., 2026). Precise instructions for data download and preformatting to enter SpaceBlocks can be found in Zenodo (doi: 10.5281/zenodo.23378236).

## Supplementary Figures

**Supplementary Figure 1.**
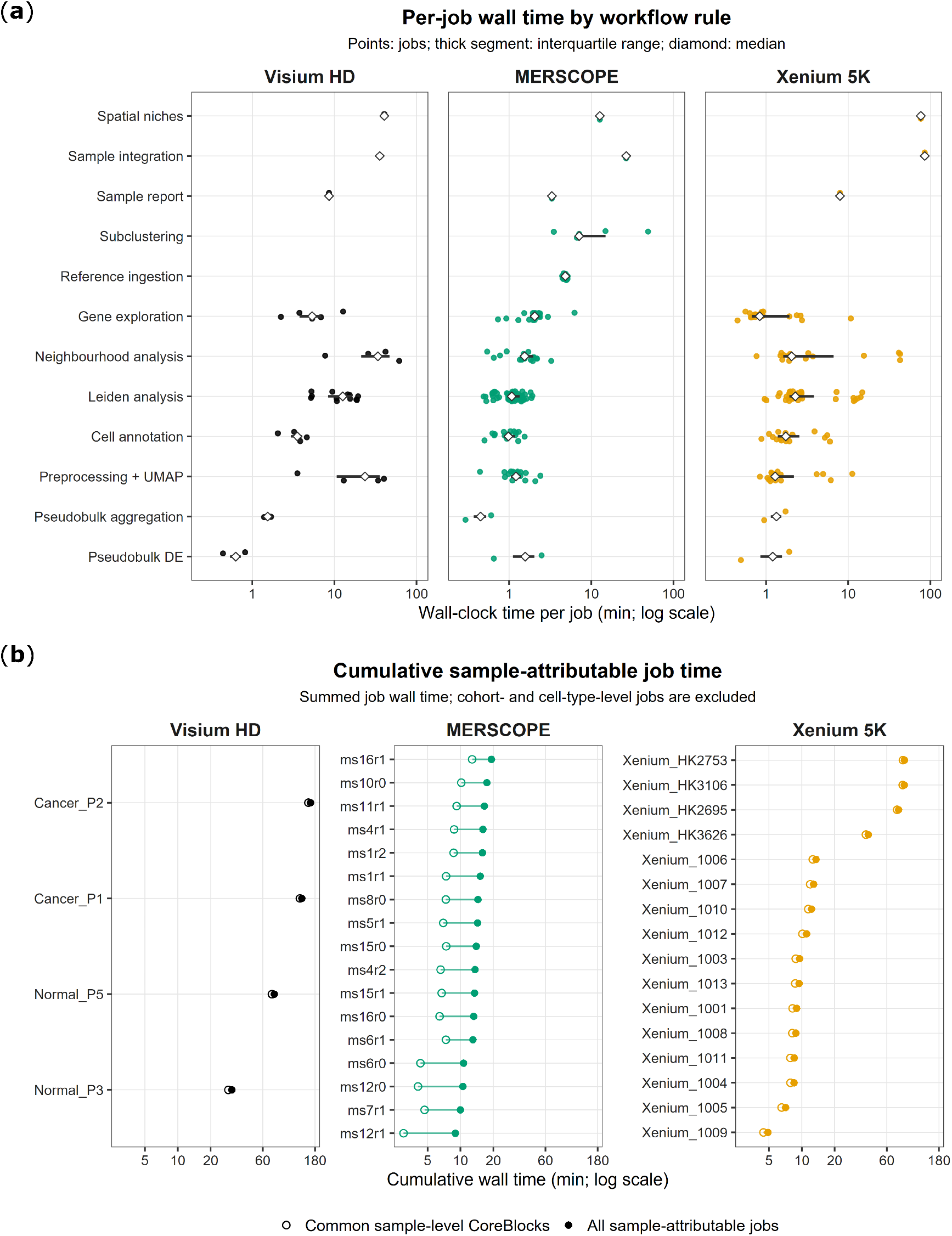
Extended benchmarking per Snakemake rule and technology. (a) Per-job wall time by workflow rule across Visium HD, MERSCOPE and Xenium 5K samples. (b) Cumulative sample-attributable job time.

**Supplementary Figure 2.**
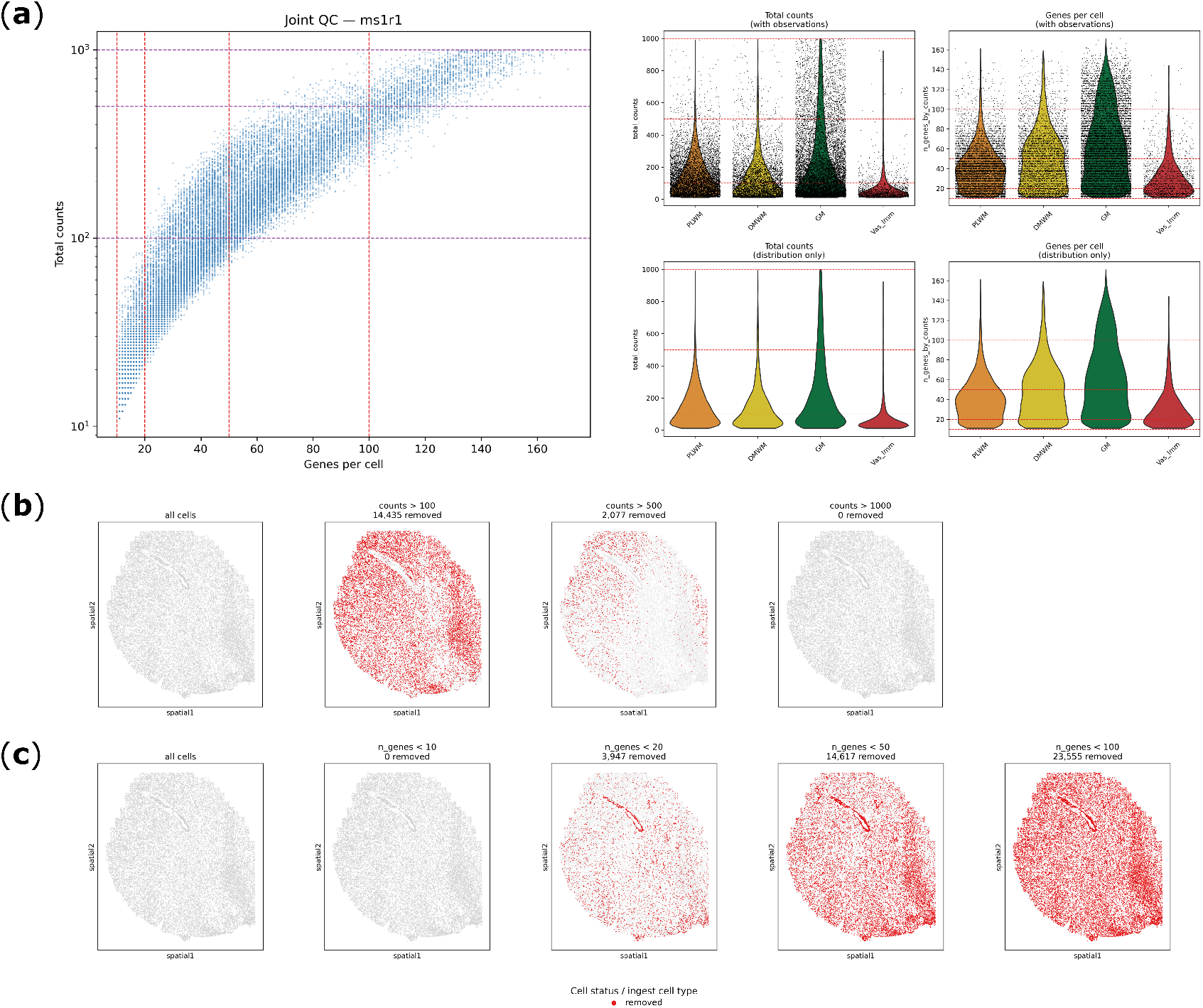
Quality control sweep diagnostic plots from the MERSCOPE dataset. Quality control diagnosis is exemplified with the ms1r1 sample. (a) Scatter plot (left) showing potential total raw counts (100, 500, 1000) and number of genes per cell (10, 20, 50, 100) thresholds with dashed lines on the ms1r1 sample. Violin plots (right) showing the same thresholds with (top) and without (bottom) individual cell observations. Violin plots are split by the annotated niches from Feng, et al. 2025. (b, c) Spatial plots indicating the number and spatial coordinates of cells that would be removed (red) and cells that would be preserved (grey) in the ms1r1 sample if the thresholds were applied for downstream analyses.

**Supplementary Figure 3.**
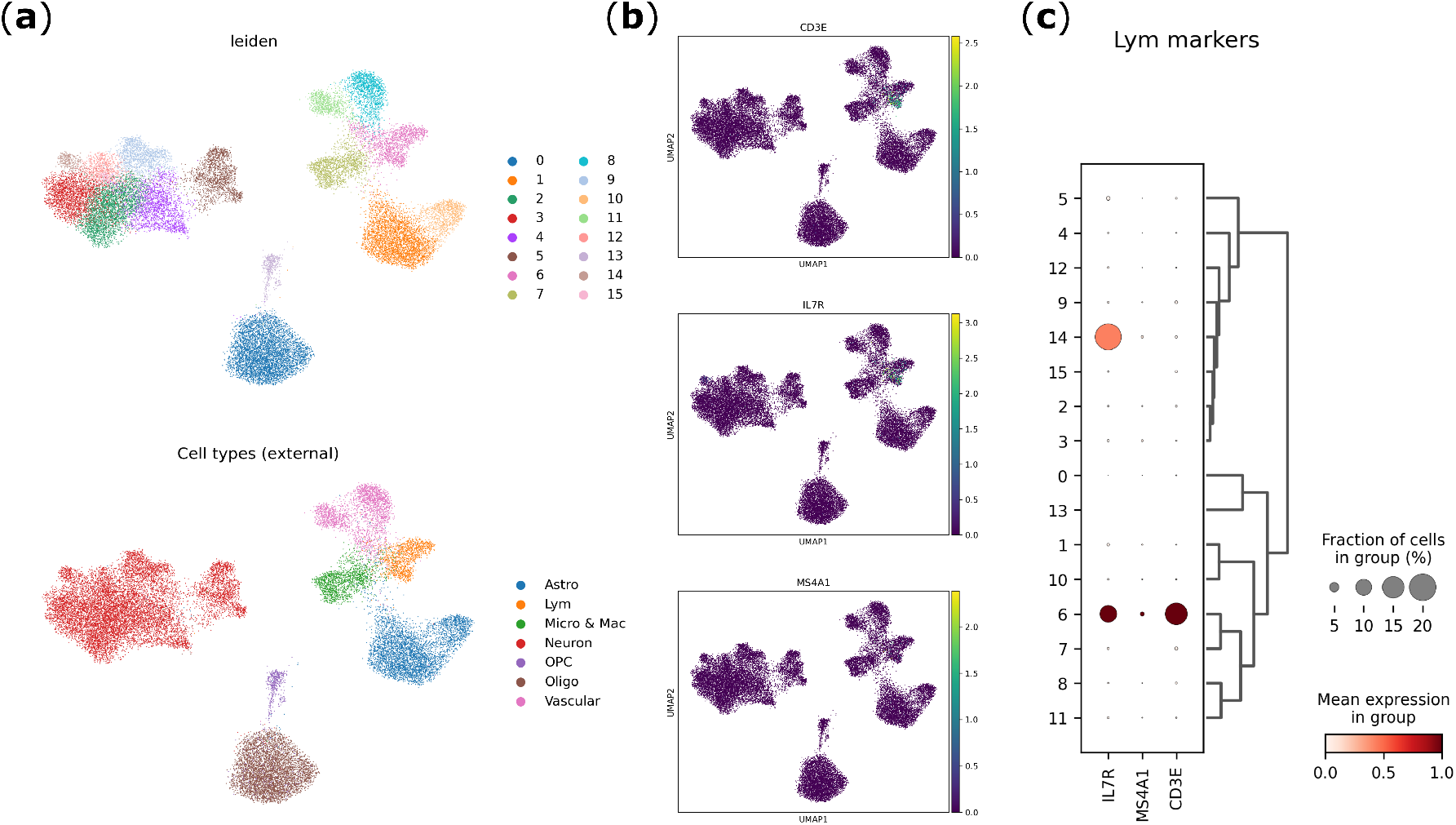
SpaceBlocks identification of heterogeneous lymphocyte marker gene expression in the MERSCOPE dataset. Sample ms1r1 from the MERSCOPE reference is used as an exemplification. (a) Comparison between SpaceBlocks Leiden clustering at 0.8 resolution (top panel) and Feng, et al. cell type annotation (bottom panel). (b) UMAP plots showcasing the expression of lymphocyte markers (CD3E, IL7R, MS4A1) described in the MERSCOPE original publication. (c) Dot plot showing the expression of lymphocyte markers in the SpaceBlocks identified clusters. Lymphocyte markers were only identified in ∼20% of cells in cluster 6, which approximately corresponds to the MERSCOPE dataset Lymphocyte annotated cells. As shown in the UMAPs, the expression was not homogeneously detected across the corresponding Lymphocyte annotated cells.

**Supplementary Figure 4.**
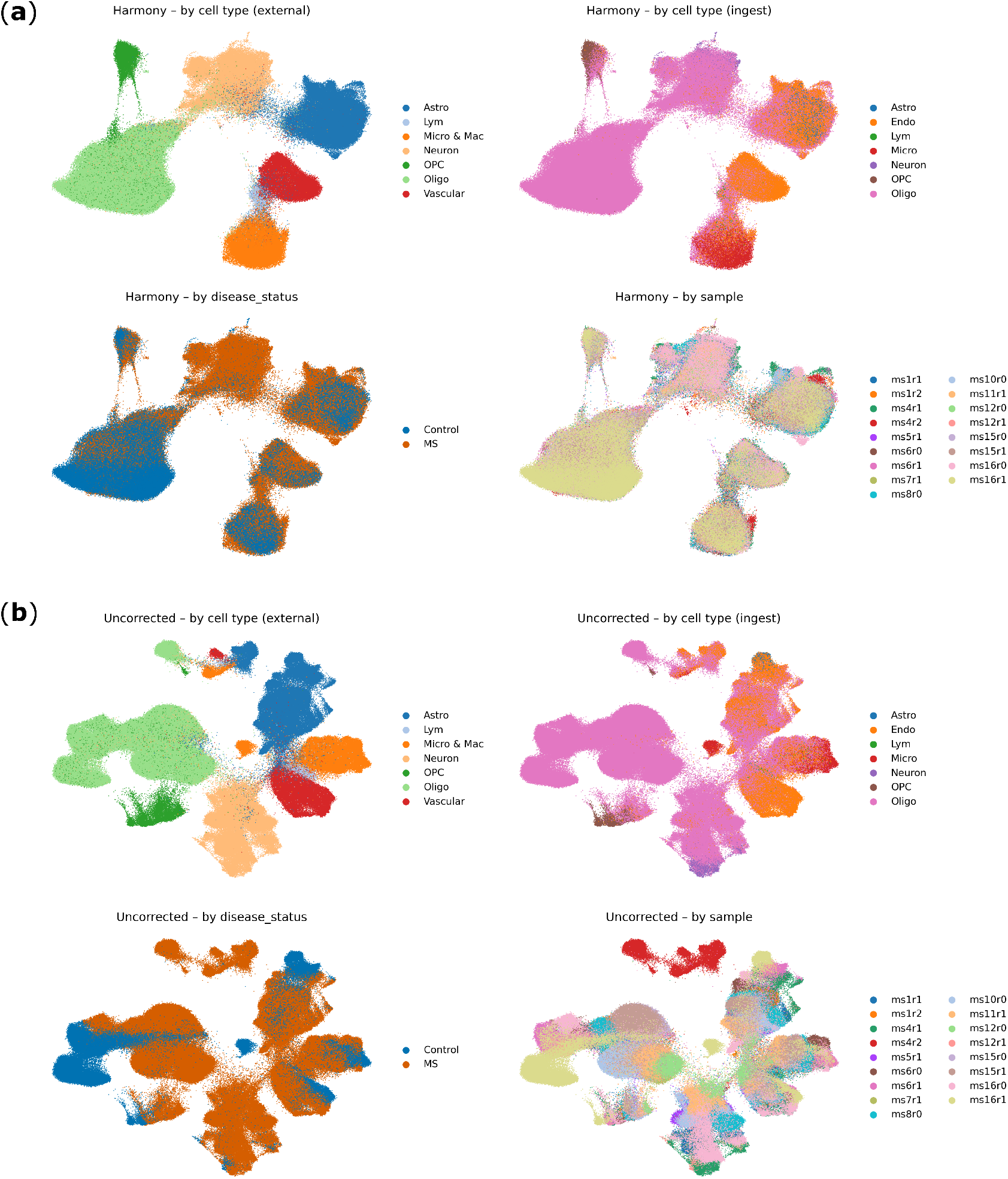
SpaceBlocks batch corrected and uncorrected UMAPs from the MERSCOPE dataset. Dataset from Feng, et al., containing 17 samples with healthy white matter (n = 3) and multiple sclerosis lesional tissue (n = 14). (a) UMAPs displaying the MERSCOPE dataset harmony integration, corrected by sample. UMAPs were coloured by different variables. From top to bottom and from left to right: original cell type annotation (Feng, et al., 2025); automatic annotation from *scanpy*.*ingest*, using the single cell dataset from Feng, et al. as a reference; disease status; and individual samples. (b) UMAPs displaying the MERSCOPE dataset without batch correction. The panels follow the same order as in (a) to ease visual comparison. MS: multiple sclerosis; Astro: astrocytes; Lym: lymphocytes; Micro & Mac: microglia and macrophages; OPC: oligodencrocyte precursor cells; Oligo: oligodendrocytes.

**Supplementary Figure 5.**
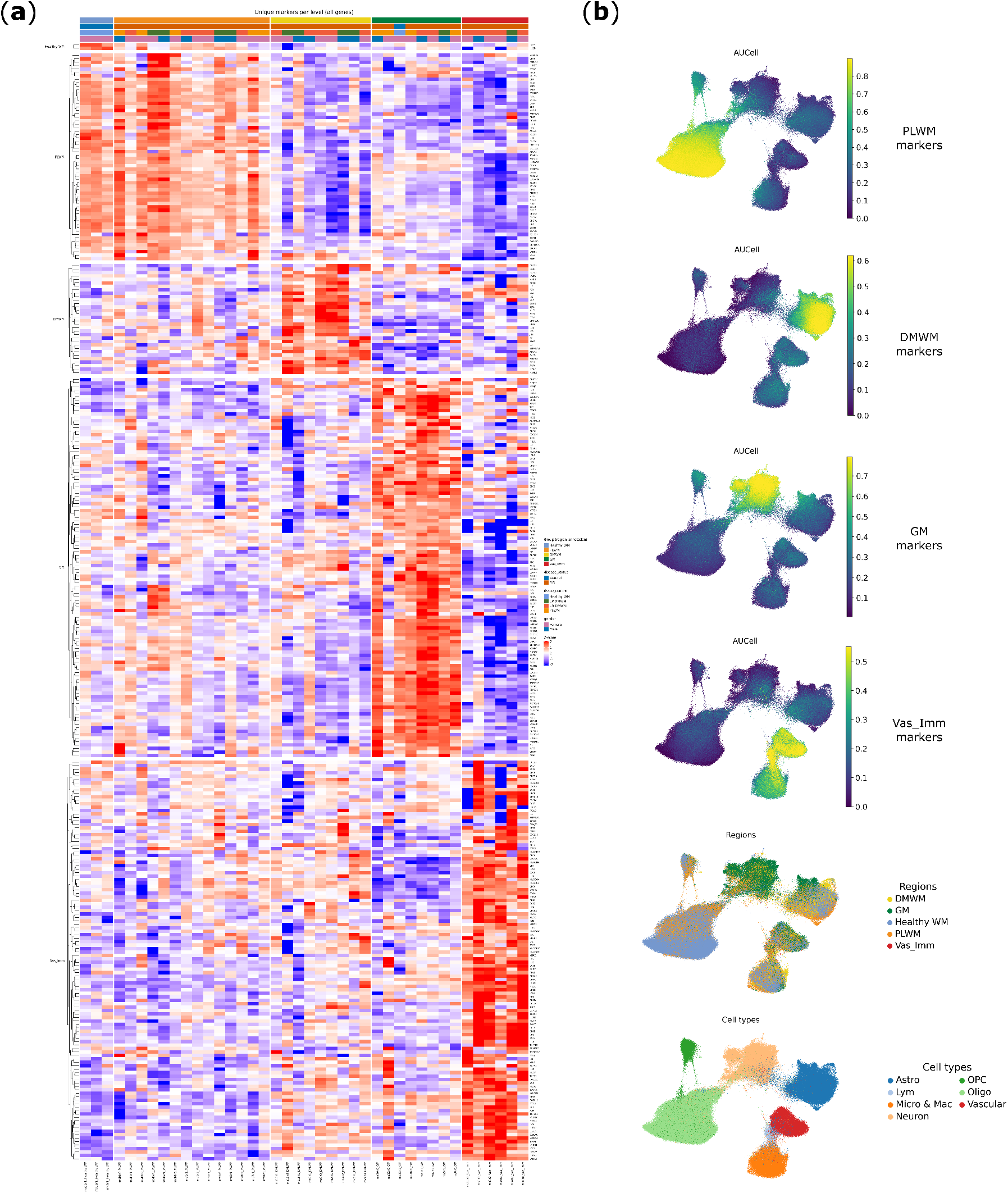
SpaceBlocks identifies differentially expressed genes between BANKSY domains in the MERSCOPE dataset. (a) Heatmap showcasing domain-specific upregulated genes from one-vs-rest tests (Benjamini-Hochberg p-adj < 0.05 and |log2FC| > 0.25) from the SpaceBlocks pseudobulk rules. Heatmap top and row annotations indicate the BANKSY domain where genes were differentially expressed. (b) UMAPs indicating AUCell enrichment from the gene sets identified in (a), calculated with SpaceBlocks exploration rules. Bottom UMAPs display the BANKSY domains and cell type annotations by Feng, et al. DMWM: demyelinated white matter; GM: grey matter; Healthy WM: healthy white matter; PLWM: perilesional white matter; Vas_imm: vascular-immune enriched area.

**Supplementary Figure 6.**
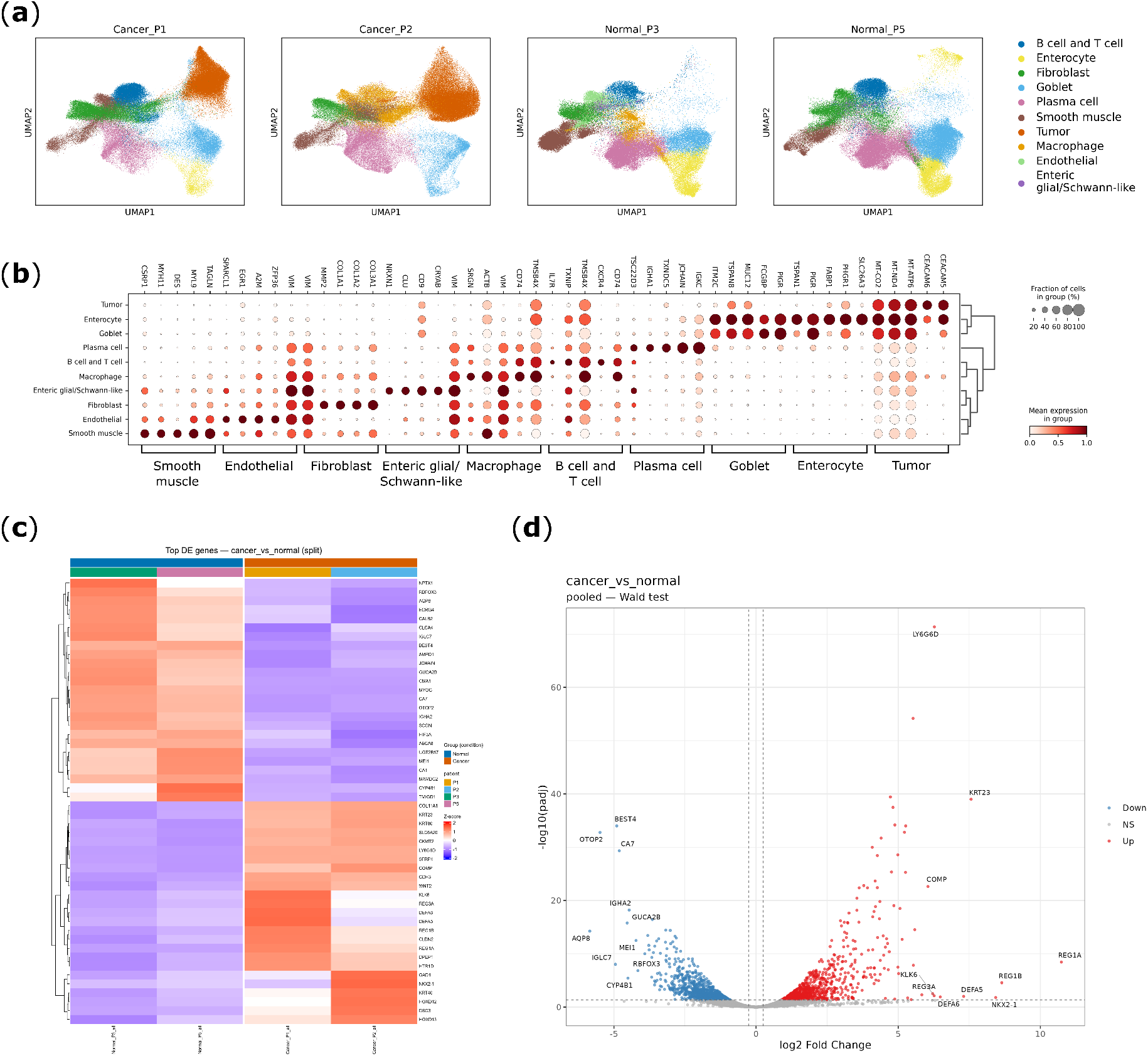
SpaceBlocks reprocessing of the 10x Genomics colon cancer dataset. Publicly available Visium HD dataset containing 2 healthy and 2 malignant colorectal cancer samples. (a) UMAPs per sample showing annotated cell types from Leiden clustering resolution 0.8. (b) Top 5 marker genes expressed by the 10 different cell types identified. (c) Heatmap displaying the top 25 marker genes with the highest and lowest fold change values between healthy and cancer conditions. (d) Volcano plot showing the differentially expressed genes between healthy and cancer conditions.

## Notes

https://github.com/cbib/SpaceBlocks

https://zenodo.org/records/23378236

### Competing Interest Statement

The authors have declared no competing interest.

